# From fluke to fragment: a multifaceted method for molecular sex identification and mitochondrial haplotyping from environmental DNA samples

**DOI:** 10.64898/2026.04.30.719183

**Authors:** Lauren Kelly Rodriguez, Sandra Schallhart, Philipp Hobmeier, Thomas Curran, Sergi Pérez-Jorge, Rui Prieto, Cláudia Oliveira, Mónica A. Silva, Bettina Thalinger

## Abstract

1. Environmental DNA (eDNA) analyses have become a powerful tool for non-invasive biodiversity monitoring, yet the applicability of population genetic approaches to environmental samples remains largely unexplored. Even when genetic traces originate from a single individual, low target DNA concentrations and amplification or sequencing artefacts can compromise downstream genetic inferences. Here, we present a novel approach for obtaining demographic insights and lineage-level mitogenomic information from aquatic eDNA samples collected near vertebrate individuals.
2. Paired eDNA and tissue samples were collected during sperm whale (*Physeter macrocephalus*) encounters in the Azores. Samples were screened for the presence of vertebrate eDNA and analyzed with a novel molecular sex identification assay. Additionally, long-range PCR was used to amplify up to five mitochondrial DNA fragments (∼3-4k bp) before subsequent sequencing on an Oxford Nanopore Technologies platform. A stringent three-tier filtering framework capable of identifying true mitogenomic variation across eDNA samples was developed for maximum recovery of genetic diversity at the haplogroup level. By benchmarking eDNA samples via their paired tissues, parameter values were optimized to maximize concordance and minimize spurious variant calls.
3. Sexing was successful for 50% of eDNA samples, with 96% concordance to paired tissues, and marine vertebrate DNA concentration significantly predicted sexing success. Further, Medaka polishing produced high identity mitochondrial consensus sequences (>16 kb) from eDNA samples. Across filtering regimes in the framework, curated SNP panels comprising up to 453 high-confidence mitochondrial SNPs resolved 19 haplogroups, with 93% concordance between eDNA and tissue samples. An intermediate bioinformatics filtering strategy maximized biologically accurate haplogroup recovery while minimizing sequencing artefacts, providing the most reliable lineage-level inferences.
4. This integrative approach demonstrates that targeted nuclear assays combined with long-range mitochondrial sequencing can recover individual-level genetic information from aquatic eDNA. By defining analytical thresholds governing success, the framework advances non-invasive genetic monitoring of populations via eDNA and enables population-level monitoring and conservation of endangered and genetically-vulnerable species.

## 1. INTRODUCTION

Environmental DNA (eDNA) analysis has rapidly emerged as a transformative approach in ecological monitoring and conservation biology. By capturing trace genetic material shed by organisms into their environment via skin cells, mucus, feces, etc., eDNA enables species detection without direct observation or disturbance to individuals (Bohmann et al., 2014). Most eDNA applications to date have focused on species-level detection, either through targeted assays or through metabarcoding approaches that enable the analysis of community composition from environmental samples (Beng & Corlett, 2020). However, there is growing interest in extending eDNA analyses–particularly for marine vertebrates of conservation concern–to population genetics applications, which would allow non-invasive estimates of genetic diversity, relatedness, sex ratio, population structure, and gene flow (Adams et al., 2019; Yates et al., 2023). Such information is central to conservation biology, as genetic variation influences adaptive potential and population viability under environmental pressures (Willi et al., 2021; Lowe & Allendorf, 2010). The earliest demonstrations of population genetic inferences from eDNA relied on mitochondrial haplotyping, and studies have proven that mtDNA haplotypes derived from seawater can match those obtained from tissue of marine megafauna, illustrating the feasibility of familial-level detection from eDNA of rare species (Jensen et al., 2021; Parsons et al., 2018; Székely et al., 2021). However, the reliance on short-barcode loci (300-500 bp) across these applications constrains eDNA-based studies to reduced resolution across the mitogenome.

More recently, long-read sequencing techniques have been validated for their potential to recover entire vertebrate mitogenomes from eDNA samples (Bista & Lino, 2026; Matthews et al., 2026; Mizuno et al., 2025). However, long DNA fragments of vertebrate taxa are only a fraction of the total amount of eDNA retained in samples and degrade faster than shorter fragments, thus reducing the potential for successful eDNA-based population genetic analyses (Gómez-Repollés et al., 2026; Székely et al., 2021). Successful population genetic inferences from eDNA samples must therefore combine cutting-edge sequencing technology with robust laboratory and bioinformatic processing to deliver maximum information from limited amounts of target DNA. Recent advances in long-read sequencing technologies (i.e., third-generation sequencing), particularly Oxford Nanopore Technologies (ONT), have created opportunities for high-quality eDNA-based mitogenome reconstruction. Long-read platforms generate multi-kilobase reads that can span entire mitochondrial loci or long-range PCR (LR-PCR) amplicons, and improvements in ONT basecalling accuracy and error correction (e.g., Guppy super-accuracy basecalling and Medaka neural network polishing) now support reliable consensus sequence generation, even from environmental extracts (Lee et al., 2021; Zhang (张天缘) et al., 2025). These advances have enabled successful mitogenome assembly from eDNA in controlled systems (Mizuno et al., 2025), suggesting the feasibility of extending this approach to the field. In this context, mitochondrial SNP (Single Nucleotide Polymorphism) panels offer a powerful and practical tool for resolving haplotypes from long-read data (Cornelis et al., 2017; Huang et al., 2023; Salas & Amigo, 2010). SNP-based haplotyping enables uniform comparisons across partially reconstructed mitogenomes, accounts for potentially non-uniform coverage across fragments, and mitigates the influence of local alignment artefacts, an important consideration when working with eDNA samples and LR-PCR products (Diroma et al., 2021).

Nuclear DNA, though less prevalent than mtDNA, remains essential for demographic inference. Sex-linked nuclear loci provide a direct insight into sex ratios, dispersal patterns, reproductive potential, and social structure, especially for vertebrate species (Korpelainen, 1990; Liu et al., 2024). Standard assays for molecular sexing target regions on sex-chromosomes are, in most cases, designed to deliver two distinct amplifications at different fragment length for the heterogametic sex, and one amplification for the homogametic sex (Morinha et al., 2012; Reddy et al., 1997; Curtis et al., 2007). Although these assays have been well established for tissue samples and feces (Thalinger et al., 2018), their application to water-derived eDNA has rarely been attempted and many of the so-far published assays are not taxon-specific and thus have limited potential for delivering reliable results from eDNA samples. Nevertheless, studies in terrestrial species (Barber-Meyer et al., 2020; Monge et al., 2020) and the most recent first application to cetacean eDNA (C. V. Robinson et al., 2025) suggest that sex-linked nuclear loci can persist in aquatic environments at detectable levels, providing another layer of population genetic information that is obtainable for target species from field-collected eDNA samples.

Cetaceans represent a critical test case for advancing eDNA-based population genetics. They are ecologically influential top predators, yet many species remain threatened by historical exploitation and ongoing anthropogenic pressures (Azzellino et al., 2017; Simmonds & Isaac, 2007). Traditional genetic monitoring in cetaceans has relied on tissue biopsies, stranded individuals, or opportunistic sampling, approaches that are powerful yet logistically difficult, spatially limited, and sometimes disruptive. eDNA offers a compelling non-invasive alternative, but reliable recovery of population-level markers (notably nuclear loci) remains technically challenging due to low DNA concentrations and degradation (T. S. Jo et al., 2022; Yao et al., 2022). Sperm whales (*Physeter macrocephalus*) are especially challenging to sample because of their deep diving behavior and extensive spatial ranges (Eguiguren et al., 2025; Whitehead, 2002). The species exhibits matrilineal social structure, limited mitochondrial diversity, and regionally-variable male dispersal, making it an informative model for testing whether eDNA can recover both mitochondrial lineages and nuclear sex markers (Alexander et al., 2016; Morin et al., 2018). The Azores archipelago, where both resident units and transient males occur, and long-term monitoring efforts have provided a baseline framework of regional matrilineal populations, provides an ideal setting for validating these eDNA method explorations (Pinela et al., 2009; Silva et al., 2014).

This study provides a novel method for obtaining population genetic information on individual vertebrate specimens via eDNA samples by integrating molecular sexing with targeted analysis of mitochondrial variants derived from long-range PCR and ONT sequencing. This approach is validated using paired eDNA and tissue samples from sperm whales as a real-world benchmark. The developed workflow encompasses targeted amplification of multi-kilobase mitochondrial fragments, super-accuracy Nanopore consensus polishing, multi-tier SNP filtering, and haplotype assignment to optimize the recovery of genetic variability from eDNA samples. By successfully demonstrating that aquatic eDNA can recover mitochondrial lineages in accordance with paired tissue samples, and that nuclear sex markers can be amplified from the same extracts, the framework provides a robust methodological foundation for extending eDNA analyses beyond species detection under real-world conditions with limited amounts of high-quality target DNA. This scalable approach for eDNA-based lineage-level (and potentially individual-level) inference in vertebrates enables non-invasive monitoring of demographic structure, mitochondrial diversity, and population connectivity without necessitating continuous sampling of reference tissues.

## 2. MATERIALS AND METHODS

### 2.1 Permits & ethical concerns

Skin biopsy sampling of sperm whales was conducted under permit LMAS-DRPM/2023/02 (CCIR/08/2023/DRCT), issued by the Environment Directorate of the Regional Government of the Azores. Biopsies were collected by trained researchers using a licensed remote biopsy system and in accordance with established best-practice guidelines for cetacean sampling. Vessel approaches were conducted from behind at controlled speed, never positioning the vessel in front of animals or groups, and sampling was performed only under suitable sea conditions to minimize disturbance. Water samples were collected non-invasively and did not require a separate collection permit under regional regulations. The transfer of genetic material between the University of the Azores and the University of Innsbruck complied with CITES (Council Regulation (EC) No 338/97, Article 7(4)) and EU Regulation (EU) No 511/2014 implementing the Nagoya Protocol. All necessary permits for collection, transfer, and analysis of genetic materials were obtained in accordance with national and international regulations.

### 2.2 Field sampling

eDNA samples (30 L) from individual sperm whale flukeprints were collected together with paired tissue samples during dedicated sperm whale monitoring surveys in summer 2023 around the islands of the Azores, Portugal (for details on field sampling, see S1 & Supporting Fig. 1).

### 2.3 Lysis, extraction, and vertebrate DNA screening

All molecular work was performed in laboratory spaces dedicated to eDNA work with separate pre- and post-PCR rooms. The laboratory workflow is summarized in Fig. 1 and detailed protocols are in S2-8. DNA extracts were primarily screened for vertebrate DNA using the MarVer3 assay by Valsecchi et al. (2020) (S4, Supporting Table 1). Successful amplification of vertebrate DNA was assessed via fluorescent signal at 245 bp. This was used to identify eDNA samples carried into subsequent molecular workflows.

**Figure 1.**
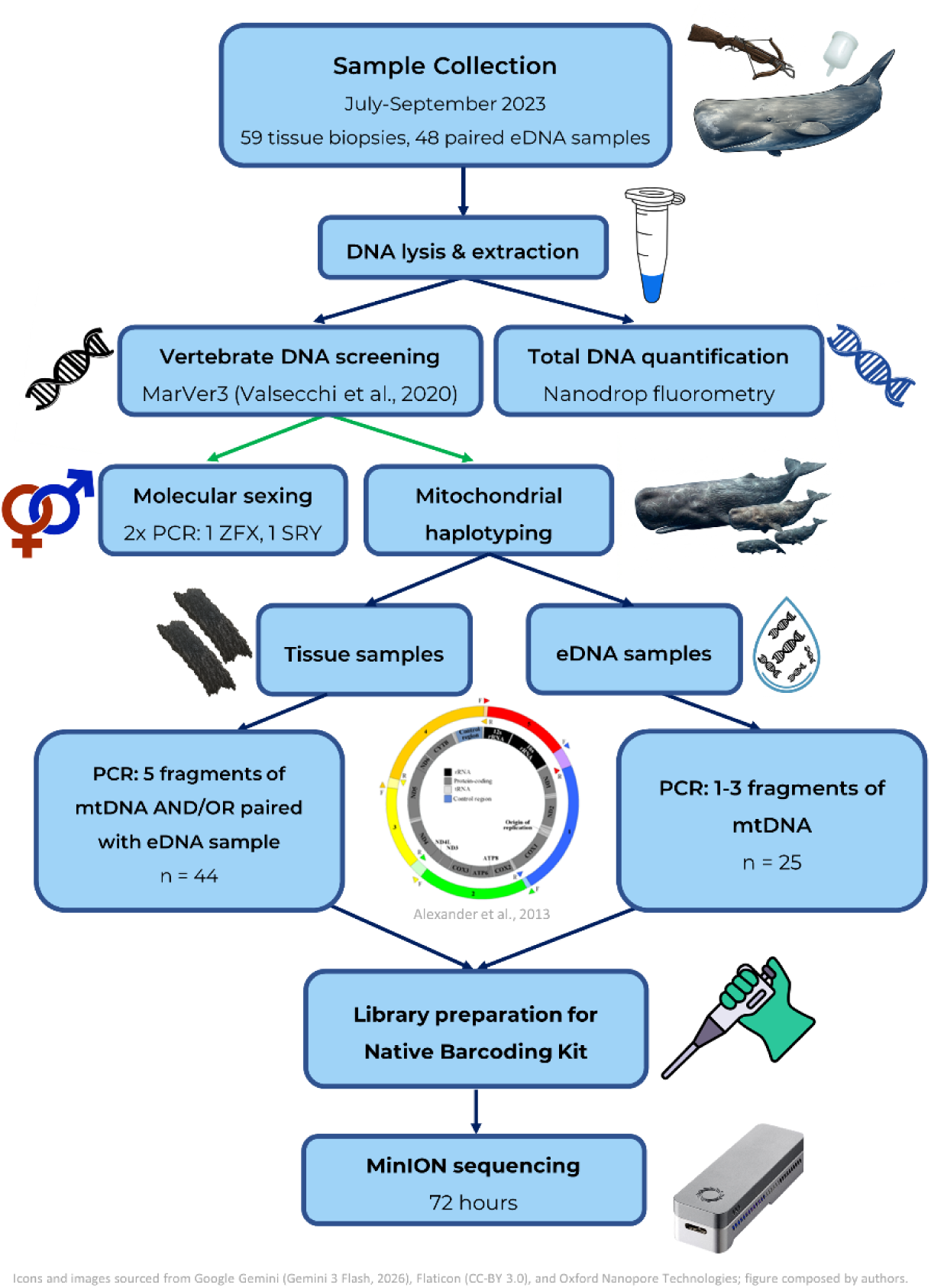
Overview of the laboratory workflow for sperm whale eDNA and tissue-based molecular analyses. Samples were collected in the Azores, and molecular processing was carried out at the University of Innsbruck.

### 2.4 Molecular sexing

Separate PCR assays were designed targeting the sex-determining region Y gene (SRY) and the zinc finger protein X-linked gene (ZFX), enabling amplification across all samples with the ZFX assay and detection of male individuals via the SRY assay (Supporting Table 1). After laboratory optimization, two subsequent PCRs were carried out (S5). Amplicon presence was assessed after the second PCR via capillary electrophoresis using a QIAxcel Advanced system (QIAGEN, Hilden, Germany) with standard DNA screening settings. All SRY and ZFX amplicons generated from eDNA samples, as well as a subset of tissue-derived amplicons, were subjected to Sanger sequencing and confirmed the amplification of sperm whale SRY and ZFX amplicons for all samples.

To evaluate the influence of marine vertebrate DNA concentration on eDNA sexing success, a binomial Generalized Linear Model (GLM) with a logit link function was fitted by maximum likelihood:

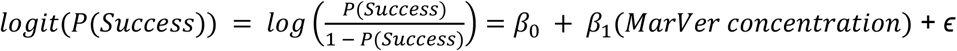

in which P(Success) represents the probability of successful amplification of sex-specific markers in an eDNA sample. *β*_0_ represents the intercept and *β*_1_ represents the change in log-odds of successful sex identification per unit increase in marine vertebrate DNA concentration.

### 2.5 Long-range mitochondrial PCR

Long-range PCR (LR-PCR) was used to amplify five overlapping fragments (3.0-4.4 kb each) spanning the mitochondrial genome of the target species, following Alexander et al. (2013) (S6, Supporting Table 1). Tissue-derived extracts were amplified for all five fragments. Three fragments maximizing the recovery of variable positions within the COX2–CYTB–control region block, containing most known mitochondrial diversity in sperm whales (Morin et al., 2018), were selected for eDNA amplification. Successful amplification was defined as recovery of at least one of the five fragments via agarose gel electrophoresis from a sample. PCR products were pooled for Nanopore sequencing by sample (eDNA or tissue). Pools from PCR fragments were quantified using a Qubit dsDNA High Sensitivity Assay (Thermo Fisher Scientific, Waltham, USA), followed by molarity-based normalization and transfer into a single barcode plate for downstream ONT library preparation and sequencing (S7-8).

### 2.6 Bioinformatics and data analysis

For clarity, key terms used throughout bioinformatics filtering are defined below (Table 1). Raw data were basecalled and demultiplexed with Guppy using the SUP model appropriate for R10.4.1 flowcells (Wick et al., 2019), producing one FASTQ file per barcode, each corresponding to a single tissue or eDNA sample. All downstream processing incorporated three distinct and restrictive quality control (QC) tiers designed to remove low quality samples, eliminate unreliable variant calls, and retain mitochondrial SNPs that behaved consistently (Fig. 2). Tier 1 read-level filtering was performed to exclude low coverage or off-target samples while retaining moderate coverage eDNA data (Ammer-Herrmenau et al., 2021), Tier 2 SNP-level filtering incorporated mapping quality, depth, and alternative allele thresholds, Tier 3 removed non-biological variation in the dataset by removing implausible or inconsistent variant patterns across samples. To evaluate the robustness of analytical stringency on SNP recovery and haplotype assignment, three hierarchical filtering profiles (henceforth referred to as strict, moderate, and relaxed) were implemented during each QC tier. The profiles differed in depth, quality, and allele frequency thresholds applied during processing, which are subsequently denoted by brackets.

**Figure 2.**
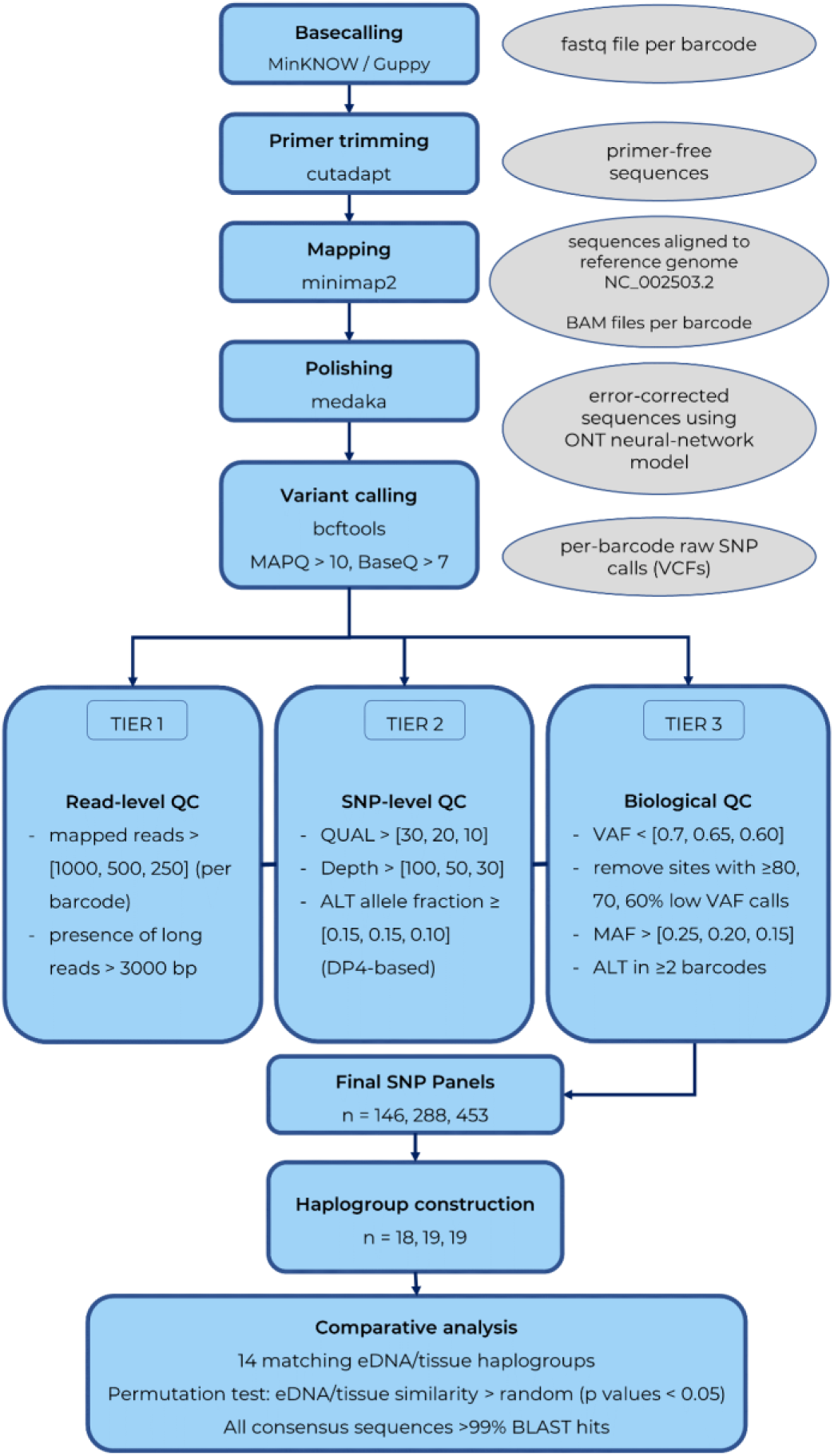
Bioinformatics workflow for mitochondrial haplotype reconstruction from Nanopore long-read sequencing of sperm whale tissue and eDNA samples. After read processing, mapping, and polishing, a three-tier quality control framework (with three profiles of filtering stringency, values per profile denoted in brackets) reduced >1,300 raw SNP positions to panels of 146 (strict profile), 288 (moderate profile), and 453 (relaxed profile) high-confidence variants suitable for comparative analyses. This panel resolved 14 matches between eDNA and tissue samples in which paired mitochondrial SNP differences were significantly lower than expected under random pairing (n = 10,000 permutations; *P* < 0.05).

**Table 1.** Definitions of terms used throughout the data processing pipeline.

| TERM | DEFINITION | SCALE |
| --- | --- | --- |
| sample | Field-collected eDNA filter or tissue biopsy | Field sample |
| pool | Combined LR-PCR amplicons derived from a single sample | Field sample |
| library | Barcoded (labeled) pool prepared for Nanopore sequencing | Nanopore library |
| read | Single DNA sequence produced by nanopore sequencer | Read level |
| basecalling | Conversion of raw ONT electrical signal into nucleotide sequence reads | Read-level |
| barcode | Unique sequencing index used to assign sequence reads to a specific library | Nanopore library |
| mapping | Alignment of sequencing reads to the reference mitogenome using minimap2 | Read level |
| profile | Predefined filtering stringency level (strict, moderate, relaxed) applied during variant calling and QC | Across samples |
| depth | Number of reads overlapping a genomic position within a sample | Position level |
| breadth of coverage | Proportion of mitogenome covered at or above a minimum depth threshold within a sample | Position level |
| VAF | Proportion of reads supporting the ALT allele at a position (ALT / (REF + ALT)) | Position level |
| variant | A SNP (as defined by the reference genome) | Position level |
| REF | Reference allele | Position level |
| ALT | Non-reference allele | Position level |
| QUAL | Phred-scaled confidence score assigned to a variant call | Position level |
| DP4 | Strand-specific allele counts (0 = RF, 1 = RR, 2 = AF, 3 = AR) | Position level |
| MAF (Minor Allele Frequency) | Across-sample allele frequency | Across samples |
| haplotype | multi-SNP genomic pattern | Across samples |
| haplogroup | A cluster of related haplotypes sharing SNP combinations | Across samples |

#### 2.6.1 Read mapping, coverage filtering, and Medaka consensus polishing (Tier 1)

After primer trimming (S9.1), reads were mapped to a mitochondrial reference genome of the target species (NC_002503.2) using minimap2 with the ONT-optimized preset. Secondary alignments were removed to ensure each read was counted once, and sorted BAM files were generated using samtools (Danecek et al., 2021). Tier 1 QC evaluated whether each barcode carried sufficient mitochondrial data to support reliable consensus generation and variant detection. The number of reads mapping to the reference mitogenome was quantified using samtools, and only barcodes with at least [1,000, 500, 250] mapped reads were retained. For each barcode passing Tier 1 QC, mapped Nanopore reads were polished with Medaka, a neural network model for correcting systematic ONT errors (Young Lee et al. 2021; Wick et al. 2023) to generate a high accuracy consensus sequence. Polishing was performed using the R10.4.1-specific SUP model r1041_e82_400bps_sup_v4.3.0. Resulting consensus FASTA files were used for variant calling and downstream analyses requiring contiguous sequence information.

#### 2.6.2 Variant calling and filtering (Tier 2)

Haplotype inference was based on curated panels of high-confidence SNPs (per profile) to minimize the influence of polishing artefacts, ensuring uniform comparison across samples and producing conservative, reproducible haplotype clusters (Jensen et al., 2021). SNPs were called separately for each barcode using bcftools applied to the Medaka-polished alignments (Genovese et al., 2024). Only biallelic SNPs were retained for downstream analyses (Quinn et al., 2024; Zhu et al., 2025). In Tier 2 QC, variants were required to have a quality score above [30, 20, 10] (Gaston et al., 2024; P. N. Robinson et al., 2017), a minimum depth of [100, 50, 30] reads (Gaston et al., 2024; Ren et al., 2024), and an alternative allele fraction of at least [0.15, 0.15, 0.10] based on strand-specific DP4 counts (Albertin et al., 2022; Diroma et al., 2021). This alternative allele threshold exceeds typical ONT and PCR error rates and remains permissive of genuine biological variation (Delahaye & Nicolas, 2021; McClinton et al., 2023). Sites failing any Tier 2 criterion were discarded. Filtered Variant Call Format (VCF) files from all barcodes were merged into a single SNP table for downstream processing.

#### 2.6.3 Position-level quality control and SNP set selection (Tier 3)

Tier 3 QC focused primarily on allele balance and consistency across samples. For each barcode/position combination, a low-confidence flag was assigned if the variant allele fraction was below [0.7, 0.65, 0.60] (Nakanishi et al., 2022). For each genomic position, the fraction of samples flagged as low-confidence was calculated. Positions where at least [60, 70, 80] percent of samples showed low-confidence values were removed, as these typically represent primer binding sites or alignment edges (S9.2; Ip et al., 2022; Smith et al., 2025). MAF was calculated for each position using high-confidence calls. Allele counts were converted to frequencies, and positions where the MAF fell below [0.25, 0.20, 0.15] were removed in order to retain SNPs that varied meaningfully across lineages (Delahaye & Nicolas, 2021; Lüth et al., 2022). To further reduce the influence of single-sample artefacts, the alternative allele at a site was required to appear as a high-confidence call in at least two different barcodes. The sites that passed all criteria formed a final set of [146, 288, 453] high-confidence informative SNPs.

#### 2.6.4 Haplotype inference

The curated SNP matrix was converted into a DNAbin-compatible format for haplotype inference using the pegas haplotype function (Paradis, 2025). Haplotypes were grouped into haplogroups for downstream analyses based on shared diagnostic mutations to avoid underrepresenting familial groups due to sequencing artefacts (Hammer & Zegura, 2002). To evaluate the sensitivity of haplogroup assignments to clustering resolution, haplotype grouping was optimized at 20 (S9.3, Supporting Fig. 2).

For barcodes in which the tissue and eDNA sample pair were retained, pairwise genetic distances were calculated on the data subset using the dist.dna in ape, computing mismatch counts across the retained SNP positions while excluding missing data (Paradis & Schliep, 2019). Concordance between pairs was based on callable positions only and defined as 1 - mismatches/called SNPs. Pairs were categorized as high-confidence matches (≥95% concordance), ambiguous (80-95% concordance), or non-matches (>20% mismatch rate). To statistically evaluate whether paired eDNA/tissue samples were more similar than expected under random association, a permutation test was conducted using the SNP distance matrix (Tran et al., 2025). A null distribution of mean distances was generated using 10,000 permutations (in R), and random eDNA/tissue pairings followed by vectorized distance extraction using purrr (MacLean, 2023; Mailund, 2022). Depth and coverage visualizations (coverage decay curves, per-position depth profiles, and SNP-level heatmaps) were generated using ggplot2, tidyverse, and dplyr (Siddiqui, 2025; Wickham et al., 2024). ChatGPT (OpenAI, GPT-5.1) was used for troubleshooting Nanopore BASH scripts; all code and parameter choices were manually verified and finalized by the authors. Full code is available at https://github.com/MarineBio-LKRod/nanopore-mito-pipeline. Basecalled FASTQ files and associated metadata are deposited in Figshare and will be made publicly available upon manuscript acceptance.

## 3. RESULTS

### 3.1 Molecular sex identification

A total of 59 tissue biopsies and 62 eDNA water samples were collected (n = 48 directly paired samples), in which eDNA samples matched the corresponding biopsy-derived sex 96% (Fig. 3a, S11, Supporting Fig. 3). The relationship between sexing success and marine vertebrate eDNA concentration was significant (Generalized Linear Model (GLM) with a log-link function; β = 0.37, *P* < 0.05, AIC = 52.16). Each 1 ng/µL increase in marine vertebrate DNA concentration increased the odds of successful sexing in eDNA samples by ∼45% (Fig. 3b).

**Figure 3.**
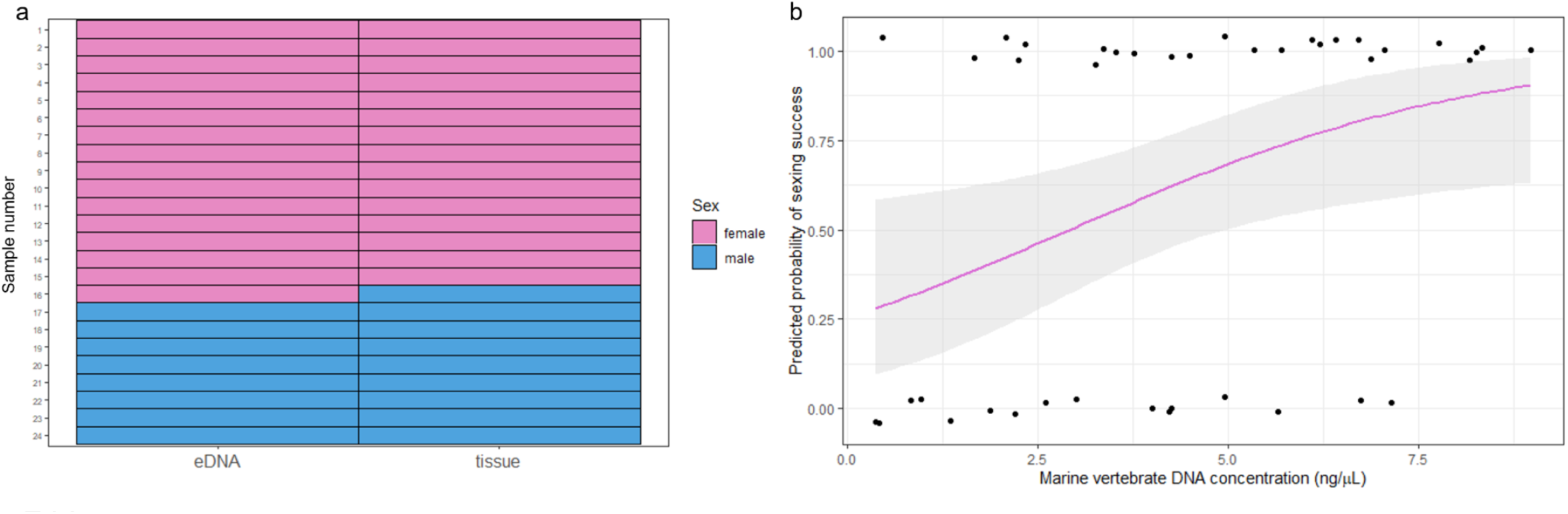
Sex-identification results from paired tissue and eDNA samples. a) Comparison of sex assignments for all matched samples, with concordant sex calls dominating the dataset and one mismatched eDNA/tissue pair. b) Logistic regression of eDNA sexing success as a function of marine vertebrate DNA concentration (ng/µL). Points represent individual eDNA samples (success = 1, failure = 0). The fitted line shows predicted probability of successful sex identification, with shaded area(s) indicating 95% confidence intervals.

### 3.2 Sequencing coverage

Across 69 samples, tissue samples consistently achieved higher mitochondrial coverage and retention under Tier 1 QC than eDNA samples, though moderate and relaxed thresholds substantially improved eDNA recovery (S10). Coverage breadth declined with increasing depth thresholds, with eDNA samples exhibiting a steeper decay than tissue samples despite all libraries passing a minimum mapped-read inclusion threshold (Fig. 4a–b; Supporting Fig. 4). In contrast to raw read count, mean mitochondrial depth after mapping emerged as a clear determinant of analytical success: samples with <60× mean depth consistently failed downstream QC tiers, whereas eDNA samples exceeding ∼350× mean depth passed all tiers and reproduced the SNP patterns of their paired tissue samples (Supporting Fig. 5). Although Medaka produced full-length (>16 kb) reference-guided consensus sequences for all eDNA libraries with >99% BLAST identity, true read-supported regions often spanned only ∼3 kb, reflecting interpolation across low-coverage regions rather than contiguous mitogenome recovery.

**Figure 4.**
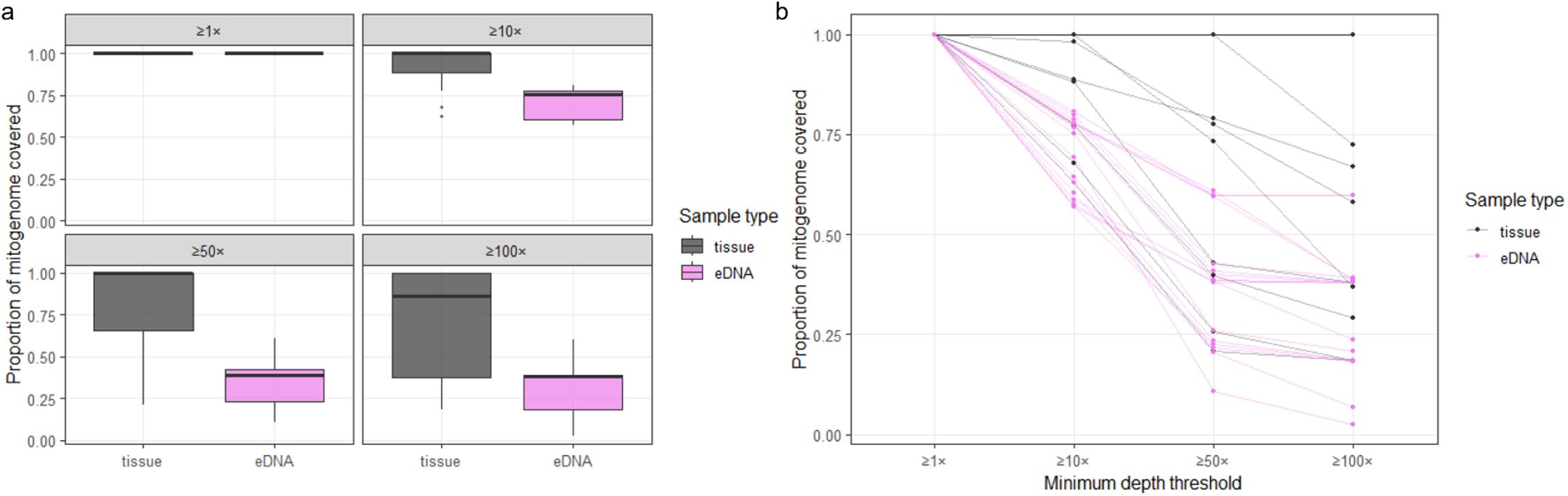
Read coverage of tissue and eDNA samples. a) Boxplots of mitogenome coverage at increasingly stringent depth thresholds for tissue and eDNA samples, illustrating contrasting coverage structures across sample types. b) Decay curves showing the proportion of the mitogenome covered at ≥1×, ≥10×, ≥50×, and ≥100× depth for each barcode, with steep decay at high thresholds in a subset of tissue samples and gradual coverage loss across eDNA samples.

### 3.3 Mitochondrial SNP panel and haplogroup resolution

Variant calling across barcodes which satisfied Tier 1 identified an initial pool of mitochondrial SNPs. Tier 2 filtering based retained 1,344, 1393, and 1449 candidate SNP positions for each profile. The subsequent Tier 3 biological filtering yielded a final curated panel of 146, 288, and 453 high-confidence SNPs (strict, moderate, relaxed, respectively), that were consistently polymorphic and supported across samples (Supporting Fig. 6). Nearly all retained SNPs fell within the COX2–CYTB–control region, with sparse representation in 12S–COX1 (Fig. 5a-c). Overall, 10, 15, and 17 paired eDNA/tissue samples were retained with four, seven, and eight high-confidence matches, under strict, moderate, and relaxed filtering profiles, respectively (Supporting Fig. 7). Although the number of distinct haplotypes increased from strict to relaxed filtering stringency, haplogroup structure remained stable across profiles (S12). Given the high number of unique haplotypes resolved across profiles, and their inherent sensitivity to minor differences in detected polymorphisms, subsequent results focus on haplogroup-level structure as the unit of population structure. Among paired samples retained under each profile, haplogroup concordance between eDNA and tissue samples was consistently high across profiles (strict: 90%; moderate: 93%; relaxed: 88%). Haplogroup structure was supported by high within-group similarity under both strict and moderate filtering (median = 1.00 and 0.99), with lower similarity observed between haplogroups (median = 0.89; 0.94, respectively). This separation was visually consistent with depth patterns across defining SNP positions (Supporting Fig. 8).

**Figure 5.**
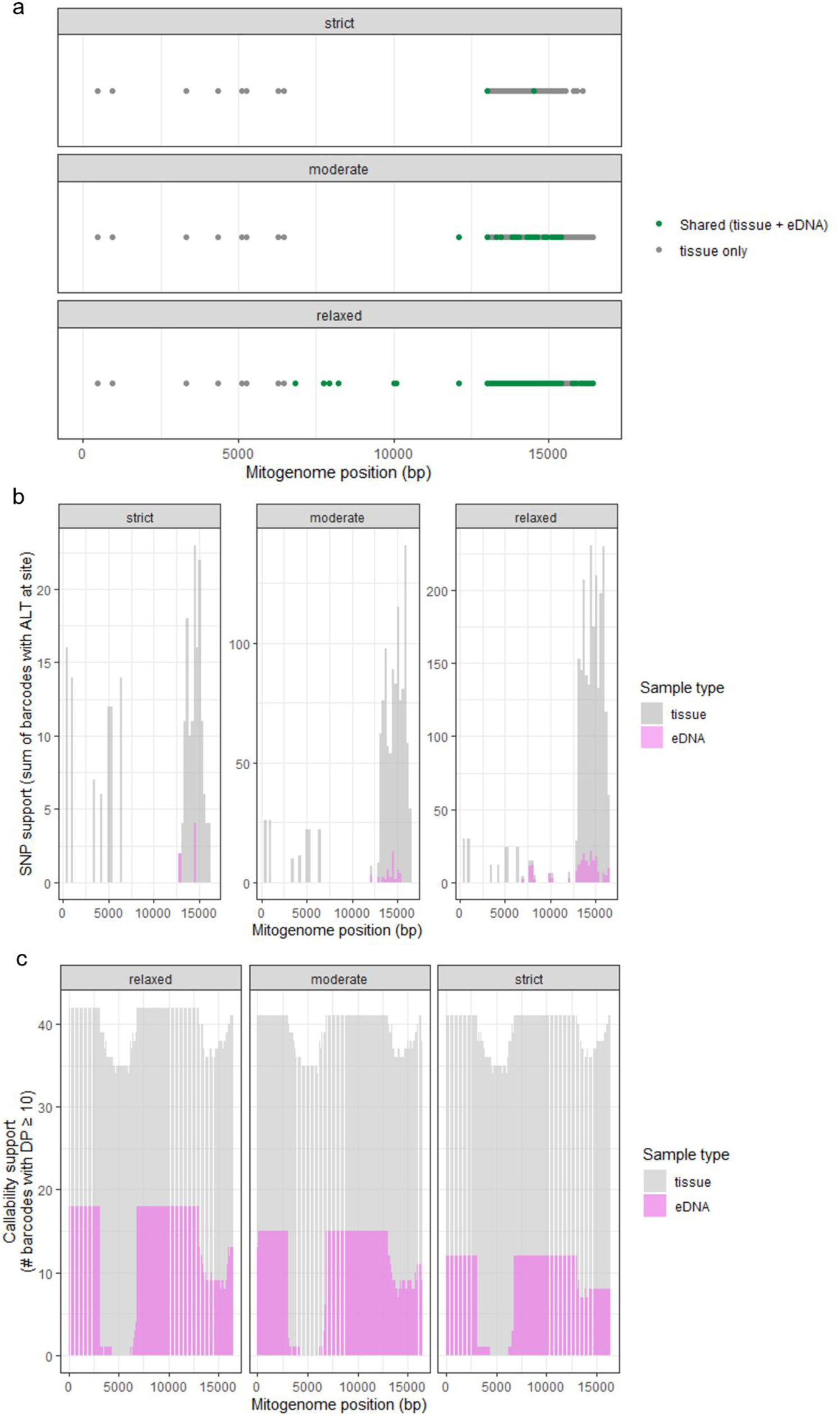
Genomic distribution and support of mitochondrial SNPs across filtering profiles. a) Positions of high-confidence SNPs along the mitogenome per profile. Points indicate SNPs observed in both tissue and eDNA samples (shared) or in tissue samples only. b) Distribution of high-confidence SNP calls across the mitogenome per profile, illustrating clustering of polymorphic sites within the COX2–CYTB–control regions. c) Proportion of callable sites (REF + ALT) across barcodes along the mitogenome for each profile.

### 3.5 Pairwise SNP distance and statistical validation

Finally, direct barcode-level comparisons demonstrated strong correspondence between paired eDNA and tissue mitochondrial SNPs (Fig. 6, S13, Supporting Table 2). Consistent with these observations, pairwise mitochondrial distances between matched eDNA/tissue pairs were significantly lower than expected under random pairing; the observed mean distance fell within the lower tail of null distributions generated from 10,000 permutations (*P* < 0.05; Supporting Fig. 9). Overall, moderate filtering retained strong haplogroup resolution while maintaining high concordance among paired samples, suggesting a balance between stringent error control and recovery of biologically informative variation.

**Figure 6.**
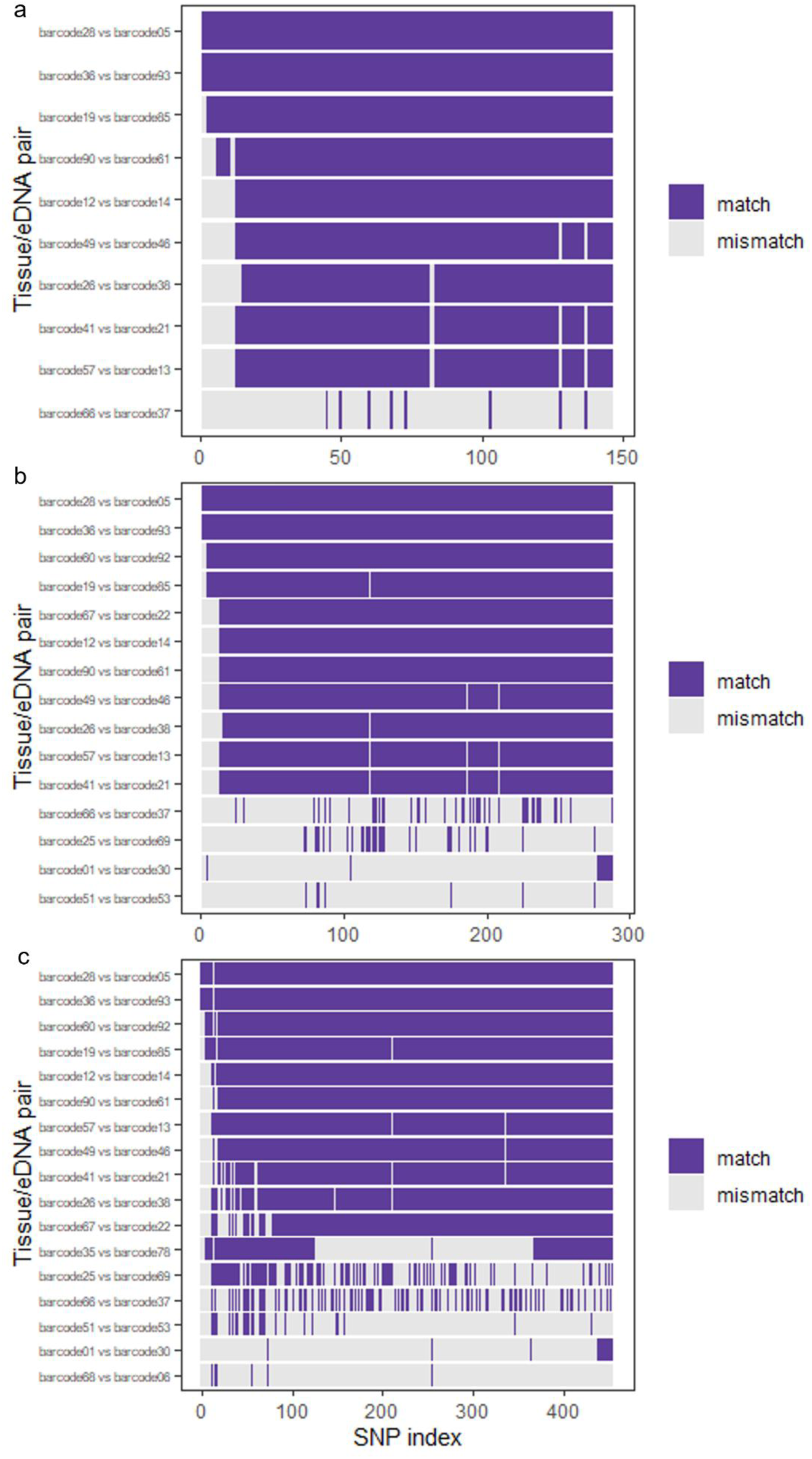
SNP-level concordance between paired tissue and eDNA samples across profiles. Panels show a) strict (n = 146 SNPs), b) moderate (n = 288 SNPs), and c) relaxed (n = 453 SNPs) with each row representing a matched eDNA/tissue pair, and each location on the x-axis corresponding to a SNP position within the respective panel.

## 4. DISCUSSION

This work presents an optimized workflow for recovering nuclear sex and mitochondrial lineage information from vertebrate individuals using aquatic eDNA. The integrated laboratory and bioinformatic pipeline enabled robust reconstruction of mitochondrial haplogroups from long-fragment eDNA sequencing data. In the presented case study of sperm whales, this approach achieved >90% concordance with paired tissue samples, providing a real-world validation benchmark. The results represent one of the first cetacean sex determinations from seawater-derived eDNA and, to our knowledge, the first reconstruction of full cetacean mitogenomes and high-resolution mitogenome SNP panels from eDNA using long-read sequencing. Together, these findings strengthen growing evidence that eDNA can recover individual- and familial-level genetic signatures even in dynamic marine environments where DNA is expected to degrade rapidly and vertebrate eDNA is comparatively rare (Adams et al., 2019; Andres et al., 2023; Székely et al., 2021).

Amplification of long mitochondrial fragments (>3 kb) from more than half of the vertebrate-positive eDNA samples indicates that long, contiguous mitochondrial DNA persists in seawater. Previous studies have amplified long fragments from eDNA in controlled environments, such as aquaria or mesocosms (Deiner et al., 2017; T. Jo et al., 2017, 2020; Mizuno et al., 2025), yet amplification success in open-ocean conditions has been considerably less predictable (Collins et al., 2018). Matthews et al. (2026) reported successful amplification of long fragments from field-collected marine eDNA in approximately 60% of their samples, which aligns exactly to our rate of recovery of long fragments. The results herein therefore confirm that long-fragment recovery is achievable *in situ* and demonstrate that the combination of LR-PCR and ONT sequencing can successfully recover extended mitochondrial reads from environmental samples. By amplifying different regions of the mitochondrial genome separately in ∼3–4 kb fragments, as opposed to the full or partial mitogenome (Matthews et al., 2026; Mizuno et al., 2025), the number of target DNA strands and thus the robustness of the method is potentially enhanced (Brandão-Dias et al., 2025; West & Deagle, 2025), providing a relatively cost-effective means to bridge the gap between short fragment eDNA barcoding and full mitochondrial genome reconstruction.

Sex identification from environmental samples clarified biological and technical constraints associated with vertebrate nuclear marker recovery. There was a significant positive correlation between sexing success and marine vertebrate DNA concentration. Hence, a marker indicating the quantity of target DNA (either through a short-fragment 16S amplicon such as the one utilized in this study or a species-specific qPCR/ddPCR assay) should be implemented as a proxy for obtaining high-quality population genetic information and enabling efficient allocation of laboratory resources. Results obtained in the present case study emphasize that extending eDNA-based analyses of vertebrate DNA to population-genetically informative nuclear markers is broadly applicable if optimized sampling and pre-processing strategies can be implemented and only one individual likely contributed the target DNA to the obtained sample (Couton et al., 2023).

Beyond overall target concentration, mitogenome coverage breadth emerged as a key determinant of analytical success across eDNA samples. Samples exhibiting broader coverage (350× and above) of the mitogenome following the reference mapping step consistently produced higher quality SNP calls and consistently matched the haplogroup delineation of their paired tissue samples across filtering profiles. This pattern, however, was not significant when assessing raw sequencing depth from environmental samples, suggesting that coverage breadth may represent a more informative quality metric than raw sequence reads in long-fragment workflows. Regions with low coverage, particularly near amplicon edges where alignment is error-prone (Zucca et al., 2016), can introduce variability that resembles true polymorphisms during mapping. Low-frequency alternative alleles at this site are consequently masked by the reference allele, reducing SNP recovery. Therefore, for rare vertebrate applications in which sample quality varies, incorporating coverage-breadth thresholds into bioinformatic pipelines will enhance confidence in lineage-level inferences.

A central concern for vertebrate population genetic inference from eDNA is whether recovered genetic diversity reflects true biological signal rather than bioinformatic artefact. Across bioinformatic profiles, haplogroup assignment for paired tissue and eDNA samples remained stable even though less stringent filtering increased the number of detected haplotypes (partly through the inclusion of additional tissue-only matrilineal lineages from unpaired samples). Higher order mitochondrial clustering and concordance between paired samples were accordingly robust to slight shifts in quality and variant allele frequency thresholds, with moderate filtering resolving the greatest number of matched pairs by balancing error removal and retention of low-coverage regions. This stability is particularly relevant for rare species applications where biological reference material and replication may be limited, and where prioritizing coarser-resolution metrics such as haplogroups can enhance reproducibility while reducing sensitivity to stochastic low-frequency alleles. Notably, the haplogroup patterns recovered here closely reflect the known matrilineal structure of sperm whales in the Azores, where social units derive from single maternal lineages but multiple lineages coexist regionally and are introduced to the area by transient males (Konrad et al., 2018).

Mitogenomic SNP panels (such as the ones generated in this analysis) are not necessarily intended to serve as a comprehensive catalog of species-wide polymorphisms, but rather as internally consistent comparative marker sets within a study (or even study region). In this context, reproducibility across samples and stability of clustering are more informative than strict concordance with previously published variant lists. Yet, if tissue samples (or other real-world biological matches) are unavailable as an empirical benchmark for this type of comparative analysis, previously-defined SNP panels can be used as a skeleton for calling alternative alleles from eDNA samples, ultimately recovering true SNPs from natural populations (Forsythe et al., 2021). In addition, sequence data generated from tissue samples (ideally from the study region) that have previously been uploaded to sequence repositories can be used as pseudo-paired mitogenomic information to train datasets and evaluate eDNA-based population genetic information. In stand-alone eDNA situations, haplogroup assignment can rely on depth-supported variants as well as replicate concordance across independent field-collected filters and/or PCRs (Goldberg et al., 2016; Shirazi et al., 2021). Additional validation strategies, including *in silico* downsampling and simulation of known haplotypes, can further constrain error rates (Barresi et al., 2025; Santorsola et al., 2016). While a general lack of tissue counterparts precludes precise estimation of false-positives and -negatives, the validation from this study serves as a proof-of-concept for broader implementation, principally across other target species with relatively low mitochondrial diversity.

## 5. CONCLUSION

The presented approach enables the identification of familial-level mitochondrial signatures and nuclear demographic markers from eDNA samples with sufficient fidelity to support lineage-level inferences. By identifying practical screening metrics (target DNA concentration), emphasizing coverage breadth as a quality indicator, demonstrating the robustness of haplogroup structure across filtering thresholds, and establishing moderate processing thresholds that retain meaningful genetic variability, this study contributes methodological advances aligned with the challenges of monitoring rare aquatic species and broadly applicable beyond the limits of the presented case study. Long-read environmental genomics, particularly when combined with quantitative screening and tiered variant filtering, offers a framework for theS non-invasive assessment of population structure, matrilineal diversity, and demographic composition. As sequencing technologies continue to improve, LR-PCR sequencing approaches may broaden the scales at which endangered marine vertebrates can be genetically monitored. Overall, these findings demonstrate the growing potential of eDNA-based analyses to extend the capabilities of conventional population genetic methods, offering new opportunities for conservation, ecological inference, and long-term monitoring of endangered species.

## Acknowledgments

This research was funded by Biodiversa+, the European Biodiversity Partnership under the 2021-2022 BiodivProtect joint call for research proposals, co-funded by the European Commission (GA No. 101052342) and with the funding organizations: Fundo Regional para a Ciência e Tecnologia (FRCT), Governo Regional dos Açores; M2.2/eWHALE/001/2023 and additionally, this research was funded in part by the Austrian Science Fund (FWF) [doi:10.55776/I6389]. MAS and RP were supported by MarAZ Researchers (01-0145-FEDER-000140) and LIFE IP AZORES NATURA – Cetáceos (LIFE17 IPE/PT/000010). CO was supported by Biodiversa+, the European Biodiversity Partnership under the 2021-2022 BiodivProtect joint call for research proposals, co-funded by the European Commission (GA No. 101052342) and the Regional Government of the Azores, through the Regional Fund for Science and Technology (FRCT), under the project EUROPAM -European Spatial-Temporal Large Scale Sea Noise Management & Passive Acoustic Monitoring of Marine Megafauna (ref. 488). CO was further supported by the Regional Government of the Azores - Portugal 2030 - European Union, and by the Regional Directorate for Science, Innovation and Development, through the PROSCIENTIA Incentive System (with the reference M1.1.C/COFUND AÇORES 2030/023/2025), under the project DETEKT - Detecting sperm whale clicks at the speed of light (ACORES2030-FEDER-01912600). SPJ was supported by NECCTON (Horizon Europe RIA GA 101081273 and the UK Research and Innovation). OKEANOS research unit received national funds through the FCT - Fundação para a Ciência e Tecnologia, I.P. by project reference UID/05634/2025 (DOI: 10.54499/UID/05634/2025), and from the Regional Directorate for Science, Innovation and Development of the Azores Government through the PROSCIENTIA Incentive System. For open access purposes, the authors have applied a CC BY public copyright license to any accepted manuscript version arising from this submission. We would like to express our gratitude to the people employed by the research institutes and all who were involved in the eDNA sampling campaign, particularly Bruno Castro. We would also like to thank Rebecca Niklas and Lea Reheis for their generous support in the laboratory. The authors declare that they have no conflicts of interest.

## Data availability statement

Raw sequencing data will be made publicly available upon acceptance.

## Author contribution statements

LKR, MAS, and BT conceived the ideas and designed methodology; MAS, CO, RP, SPJ and LKR collected and processed field-collected samples; LKR, SS, PH, and TGC performed laboratory analyses; LKR and PH analyzed the data; LKR led the writing of the manuscript, MAS and BT acquired funding, BT led project administration. All authors contributed critically to the drafts and gave final approval for publication.

## Notes

https://github.com/MarineBio-LKRod/nanopore-mito-pipeline

### Competing Interest Statement

The authors have declared no competing interest.

