## Supplementary material for "From fluke to fragment: a multifaceted method for molecular sex identification and mitochondrial haplotyping from environmental DNA samples": S1

### Supporting Methods

#### S1. Field sampling

Water samples were collected opportunistically from surface flukeprints immediately following sperm whale dives and ensuring that only one individual had transversed the water column at the flukeprint position. At each encounter, ~30 L of seawater was pumped through single-use filter capsules (Waterra, Mississauga, CA; 0.45 µm pore size, 600 $cm^2$ surface area) using a peristaltic pump (Eijkelkamp, Fiesbeek, NL). Filters were preserved with 50 mL of TES buffer (0.1 M TRIS, 10 mM EDTA, 2% sodium dodecyl sulfate; pH 8) supplemented with proteinase K (20 mg/mL, 190:1 ratio; PanReac Applichem, Barcelona, Spain) and stored at -20 °C at the [removed for review] before shipment (on ice, in styrofoam containers) to the [removed for review] for storage at -80°C until subsequent laboratory processing.

Paired biopsy samples were collected from the same sperm whale individuals, whenever possible. Biopsies were obtained by licensed personnel using a Barnett crossbow (Barnett Crossbows, Tarpon Springs, USA) with Ceta-Dart biopsy darts (CETA-DART, Copenhagen, Denmark), targeting the dorsal flank below the dorsal fin. Tissues were preserved in 95% ethanol, stored, and transported under the same conditions as eDNA samples.

#### S2. Laboratory contamination control

Work surfaces were routinely decontaminated with 10% bleach followed by 80% ethanol, and all personnel wore DNA-free laboratory clothing including disposable coveralls, gloves, face masks, and hair coverings, with gloves changed frequently. The handling of PCR reagents and reaction preparations took place in a separate room equipped with PCR-dedicated workstations that were UV-sterilized ten min prior to work. To monitor for potential contamination, negative controls generated in the field, during extraction, and PCR were processed alongside all samples (Thalinger et al., 2021).

#### S3. DNA lysis and extraction

Waterra filter capsules were processed using a precipitation-based protocol adapted from Pont et al. (2018). After filter capsules were agitated on an orbital platform shaker at 300 shakes for approximately 15 min to homogenize the lysate, the contents were transferred into prelabeled DNA-free 50 mL tubes and centrifuged at 15,000 g for 15 min at 6 °C, after which the supernatant was carefully removed, leaving roughly 15 mL of liquid above the pellet. Ethanol precipitation was performed by adding 33 mL of absolute ethanol and 1.5 mL of 3 M sodium acetate to each tube, mixing gently, and storing the tubes at −20 °C for at least 12 h.

Following precipitation, tubes were left undisturbed at room temperature for at least 30 min to facilitate pellet dissolution and followed by centrifugation at 15,000 g for 15 min at 6 °C. Supernatants were decanted carefully and tubes were individually air-dried in a closed laminar flow hood for 5 min. The resulting DNA pellet was resuspended in 720 µL of TES buffer and 20 µL Proteinase K. Lysates were incubated at 56 °C for one hour and subsequently transferred to prelabeled 2 mL PCR-clean tubes for DNA extraction.

For tissue lysis, approximately 25 mg of tissue was excised from the original skin sample using sterilized scissors, with particular care taken to primarily select skin while minimizing the inclusion of adipose tissue. To prevent cross-contamination, scissor blades and tweezers were immersed in ethanol and flame-sterilized three times between samples. For improved lysis efficiency, excised tissue fragments were further sectioned into smaller pieces and transferred to individual 1.5 mL microcentrifuge tubes. The samples were air-dried under a laminar flow hood with tube lids open, positioned 30 cm apart, for 2–3 hours until ethanol evaporated completely. If residual ethanol remained after 3 hours, the drying process was extended overnight. An empty control tube was included alongside the samples as a negative control to check for potential environmental or cross-sample contamination. During each drying step, a maximum number of 10 samples plus one control tube was kept under the laminar flow hood. After complete drying, each sample was resuspended in 180 µL TES buffer and 20 µL Proteinase K. The tubes were vortexed briefly to ensure homogeneity and subsequently centrifuged shortly. Samples were then incubated at 56°C overnight on a light shaking platformer until complete tissue lysis was achieved. After incubation, the lysates were cooled to room temperature before being stored at -20°C for further analysis.

Lysates were extracted using the BioSprint 96 platform (QIAGEN, Hilden, Germany) with the BioSprint 96 DNA Blood Kit (QIAGEN, Hilden, Germany), following the manufacturer's protocol, with elution in 100 µL TE buffer (Rodriguez & Thalinger, 2024). eDNA and tissue lysates were processed separately. Extracted DNA was quantified using a Qubit dsDNA HS Assay (Thermo Fisher Scientific, Waltham, USA).

#### S4. Vertebrate DNA screening using MarVer3 primers

PCRs were carried out in 10 µL reactions containing 5.0 µL of 2× SuperFi Master Mix (Thermo Fisher Scientific, Waltham, USA), 1.7 µL molecular-grade water, 0.3 µL recombinant Bovine Serum Albumin (reBSA, 10 mg/mL; New England Biolabs, Beverly, USA), 0.5 µL of the MarVer3 forward primer (10 µM), 0.5 µL of the MarVer3 reverse primer (10 µM), and 2.0 µL of DNA extract. Amplifications were performed using a touchdown PCR to enhance primer binding specificity (Don et al., 1991). The cycling profile consisted of an initial denaturation at 94 °C for 3 min, followed by an annealing temperature decrease from from 63 °C to 57 °C in 0.5 °C increments per cycle and followed by eight cycles at 57 °C. All cycles used a denaturation step at 94 °C (30 s) and an extension step at 72 °C (30 s), the program concluded with a final elongation at 72 °C for 5 min.

Presence of marine vertebrate DNA in each sample was then assessed by capillary electrophoresis of the PCR products (diluted 1:10 in TE buffer (pH 7.0)) via a QIAxcel Advanced capillary electrophoresis system (QIAGEN, Hilden, Germany). Successful amplification produced a distinct fluorescent band (peak) at approximately 245 bp, corresponding to the expected MarVer3 fragment, which was used to identify eDNA samples to be carried into the subsequent molecular workflow.

#### S5. Molecular sexing assay design & PCRs

Sequence alignments were generated in BioEdit 7 (Hall 2004) using the target species (sperm whale) reference sequences AB108515.2 (SRY) and AF260801.1 (ZFX, exon 4) together with corresponding human sequences. Manual inspection of these alignments was used to identify conserved sperm whale-specific regions suitable for primer binding (Grubwieser et al., 2006; R. Hughes-Stamm et al., 2011; Saito & Doi, 2021). Genomic fragments within the SRY and ZFX sequences with the following characteristics were manually identified as primer candidates: maximum mismatches to human DNA, shorter than 400 bp for efficient amplification from potentially degraded eDNA templates, minimum secondary structures, and Tm between 50 and 62°C (Grubwieser et al., 2006; R. Hughes-Stamm et al., 2011; Saito & Doi, 2021). For the SRY assay, the forward primer was adapted from Nishida et al. (2003) and paired with a newly designed reverse primer to obtain a 127 bp fragment (Supporting Table 1). To enhance specificity, Locked Nucleic Acid (LNA) bases were incorporated at key positions in both forward and reverse primers, increasing primer/template binding stringency and improving mismatch discrimination (Wengel et al., 1999; You et al., 2006). For the ZFX assay, novel primers were designed to obtain a 336 bp fragment (Supporting Table 1). Candidate primers were evaluated in Primer3 (Untergasser et al., 2012) and screened using NCBI Primer-BLAST (Ye et al., 2012) to assess melting temperatures, secondary structures, and *in silico* specificity.

Laboratory optimization included gradient PCRs using target and non-target (*Felis catus*, *Canis familiaris*, *Homo sapien* male and female, *Delphinus delphis*, *Kogia breviceps*, *Kogia sima*, and *Megaptera novaeangliae*) tissue-derived DNA to ensure the absence of non-target amplification. PCR reactions were carried out in 10 µL reaction volumes using the QIAGEN Multiplex PCR Master Mix. SRY reactions contained 5.0 µL of 2× Multiplex Master Mix, 0.5 µL of the forward primer Pm-SRY-F (10 µM), 0.5 µL of the reverse primer Pm-SRY-R (10 µM), 0.5 µL bovine serum albumin (BSA, 10 mg/mL), 3 µL of template DNA, and 0.5 µL molecular-grade water. ZFX reactions were prepared equivalently using primers Pm-ZFX-F and Pm-ZFX-R (10 µM each). All PCR plates included a molecular-grade water blank (PCR negative control) and a human male DNA extract as an amplification control. All reactions were run under the same thermocycling profile: an initial denaturation at 95 °C for 15 min, followed by 30 cycles of 94 °C for 30 s, annealing at 55 °C (SRY) or 58 °C (ZFX) for 90 s, and 72 °C for 60 s, with a final extension at 72 °C for 10 min. Following the first round of amplification, PCR products were purified using a magnetic bead-based protocol (AMPure XP, Beckman Coulter) on the BioSprint automated platform (QIAGEN). For each reaction, 9 µL of PCR product was combined with 7.2 µL of AMPure beads (0.8× ratio), transferred to the BioSprint system, washed with 80% ethanol, and eluted in 20 µL molecular-grade water (hereafter “PCR clean-up”).3 µL of the cleaned PCR product were added to the second round of PCR instead of template DNA.

#### S6. Long-range PCR chemistry and cycling parameters

All reactions were performed in 20 µL volumes using Platinum™ SuperFi™ PCR Master Mix (Thermo Fisher Scientific, Waltham, USA). Each reaction contained 10 µL of 2× SuperFi Master Mix, 0.5 µL reBSA (10 mg/mL), 0.5 µL forward primer (10 µM), 0.5 µL reverse primer (10 µM), 3 µL DNA extract, and molecular-grade water. LR-PCRs were run under the following cycling conditions: an initial denaturation at 94 °C for 3 min; 40 cycles of 94 °C for 30 s, fragment-specific annealing at 60 °C (fragments 1 and 3), 61 °C (fragments 2 and 4), or 64 °C (fragment 5) for 30 s, and 72 °C for 90 s; followed by a final extension at 72 °C for 10 min.

Each amplification was subject to a PCR clean-up and eluted in 15 µL molecular-grade water. Cleaned products were then subjected to a second round of amplification using identical reaction compositions and cycling conditions, substituting the DNA extract for 3 µL of the cleaned PCR product. A second PCR clean-up (15 µL elution) was performed to remove residual primer dimers and PCR additives.

#### S7. Nanopore library preparation

Approximately 50 fmol of each cleaned pool (i.e., amplicon libraries obtained from individual tissue or eDNA samples containing 1-5 mtDNA fragments) were used as input for end-prep in which each sample was assigned a unique barcode. Following barcode ligation, all samples were pooled into one tube and quantified via Qubit dsDNA High Sensitivity Assay.
Sequencing adapters were then ligated to the pooled barcoded library, followed by two bead clean-ups to remove unincorporated adapters and barcodes before elution in ONT Elution Buffer and quantification via Qubit fluorometry to ensure that at least 50 fmol of DNA (appropriate for 1-10 kb sequences) was available for flow-cell loading, consistent with the recommended input amounts for R10.4.1 MinION flow cells (FLO-MIN114; Oxford Nanopore Technologies, 2022; Oxford Nanopore Technologies, Littlemore, UK). Immediately before sequencing, the library was prepared for loading using ONT Sequencing Buffer and Library Beads.

#### S8. Nanopore sequencing

Sequencing libraries were prepared using the ONT Ligation Sequencing Kit V14 (SQK-LSK114; Oxford Nanopore Technologies, Littlemore, UK) in combination with the Native Barcoding Expansion 96 V14 (SQK-NBD114.96, Oxford Nanopore Technologies, Littlemore, UK). Library preparation followed the manufacturer’s recommended workflow. A primed R10.4.1 flow cell was loaded on a MinION Mk1B device. Sequencing proceeded for 72 hours under standard MinKNOW settings with on-board basecalling and demultiplexing using the NBD114.96 kit configuration.

#### S9. Bioinformatic processing

##### S9.1 Primer trimming

All raw barcoded FASTQ files were processed with Cutadapt (Martin, 2011) to remove primer sequences. LR-PCR primer sequences were supplied as trimming targets at the 5′ end of reads. Trimming required a minimum overlap of 15 bases between primer and read and allowed up to 10% mismatches to accommodate both primer mismatches and potential errors related to ONT sequencing (Baloğlu et al., 2021; Rousseau et al., 2025). Reads lacking a sufficient primer match were left unchanged.

##### S9.2 Mitogenome coverage insights

Note that because mitochondrial DNA was amplified using LR-PCR from five overlapping regions in tissue samples but only up to three regions in eDNA samples, the resulting SNP panel inherently reflected uneven recoverability across sample types. Tissue-based LR-PCR amplified the full mitogenome, but eDNA libraries consistently produced strong coverage for

COX2 through the control region, which have been widely used in previous sperm whale population genetics studies. SNPs located in the 12S, 16S, ND1, ND2, and COX1 regions were absent in eDNA samples and therefore removed during Tier 3 filtering. Conversely, positions within COX2, ATP8, ATP6, COX3, ND3, ND4L, ND4, ND5, ND6, CYTB, and the control region passed quality thresholds at higher rates across both sample types and formed the majority of informative polymorphisms retained. Thus, the final SNP panel reflects both biological variation and the distribution of reliably amplified fragments, providing a consistent set of markers that could be compared across eDNA and tissue-derived sequences.

##### S9.3 Haplogroup determination

To identify an appropriate threshold at which to restrict haplotype assignment, haplotypes were hierarchically grouped to produce "capped" haplogroup sets (maximum = 10, 20, or 30 groups). For each cap, the proportion of eDNA samples falling within haplogroups that also contained tissue samples were quantified and the threshold which represented a consistent level of resolution was retained (Supporting Fig. 2).

---

### Supporting Results

#### S10. LR-PCR amplification success and mitochondrial mapping performance

Thirty-five tissue samples produced all five LR-PCR fragments; the remaining 11 tissue samples produced 2-4 fragments. Ten eDNA samples amplified one fragment, eight amplified two, and seven amplified three. Across the 69 samples (44 tissue, 25 eDNA) that yielded mappable reads, tissue barcodes averaged 55,002 raw reads (median 48,223) and 49,835 mitochondrial-mapped reads, whereas eDNA samples averaged 13,212 raw reads (median 8,074) and 5,809 mapped reads. Mapping efficiency was high in tissue samples (mean 0.90) and moderate in eDNA samples (mean 0.44). Applying strict Tier 1 QC (≥1,000 mitochondrial-mapped reads), 41 of 44 tissue samples (93%) and 12 of 25 eDNA samples (48%) were retained. For moderate and relaxed Tier 1 QC (≥500 and ≥250 mitochondrial-mapped reads), 15 eDNA samples (60%) and 41 tissue samples (93%) as well as 18 eDNA (72%) and 42 tissue samples (95%) were retained, respectively.

#### S11. Vertebrate and molecular sex identification screening

Six eDNA field negative controls did not yield amplification of marine vertebrate DNA across three replicates. Vertebrate DNA was successfully amplified in all tissue samples and in 42 of the 56 field-collected eDNA samples. Sex identification PCRs produced clear ZFX amplification for all tissue samples and additionally, 20% of the tissues were classified as male after successful amplification with the SRY assay. Sexing was successful for 50% of paired eDNA samples.

#### S12. Mitochondrial SNP panel and haplogroup resolution

The strict filtering profile (based on the panel of 146 SNP sites) resolved 24 mitochondrial haplotypes (with 18 main haplogroups). The moderate profile (n = 288 SNP sites) resolved 38 haplotypes with 19 haplogroups. The relaxed profile (n = 453 SNPs) resolved 45 haplotypes with 19 haplogroups. Fourteen diagnostic SNPs separated the two major haplogroups under strict filtering (17 under moderate filtering, 22 under relaxed filtering). Small haplogroups exhibited coherent coverage profiles, whereas the dominant haplogroup (HG4) showed broader depth variation but consistent support across informative SNPs.

#### S13. Pairwise SNP distance

Two eDNA samples produced exact matches (0 SNP differences) to their paired tissue barcode across callable SNP panels (for both strict and moderate profiling; Fig. 6a and 6b). Four additional eDNA samples differed from their tissue counterpart by less than 10 SNPs (Supporting Table 2). Across retained pairs, median SNP distances remained low relative to panel size. Under strict filtering, the median pairwise distance between samples was 13 SNPs (IQR 3-16). Moderate filtering yielded a median distance of 15 SNPs (IQR 10-127), while relaxed filtering resulted in a median distance of 29 SNPs (IQR 15-253). When normalized by panel size, median proportional SNP differences (between pairs) corresponded to 8.9% (strict), 5.2% (moderate), and 6.4% (relaxed) of the respective panels.

#### Supporting information references

9) Rousseau, C., Henry, N., Rousvoal, S., Tanguy, G., Legeay, E., Leblanc, C., & Dittami, S. M. (2025). A Practical Comparison of Short- and Long-Read Metabarcoding Sequencing: Challenges and Solutions for Plastid Read Removal and Microbial Community

Exploration of Seaweed Samples. *Molecular Ecology Resources*, *25*(7), e14129. https://doi.org/10.1111/1755-0998.14129

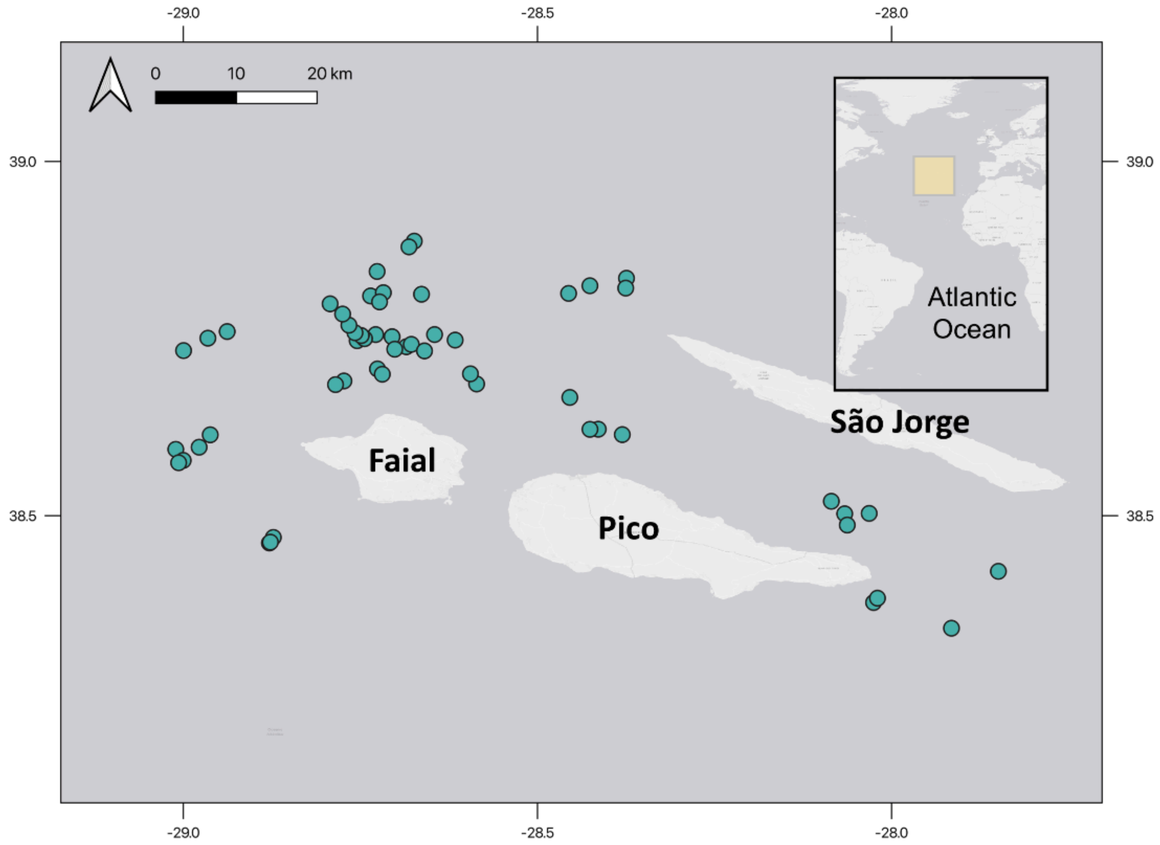


Supporting Figure 1. Sampling locations (points) of sperm whale tissue and eDNA sampling events around the islands of Faial, Pico, and São Jorge (Azores, Portugal). The inset map shows the position of the study area in the North Atlantic.

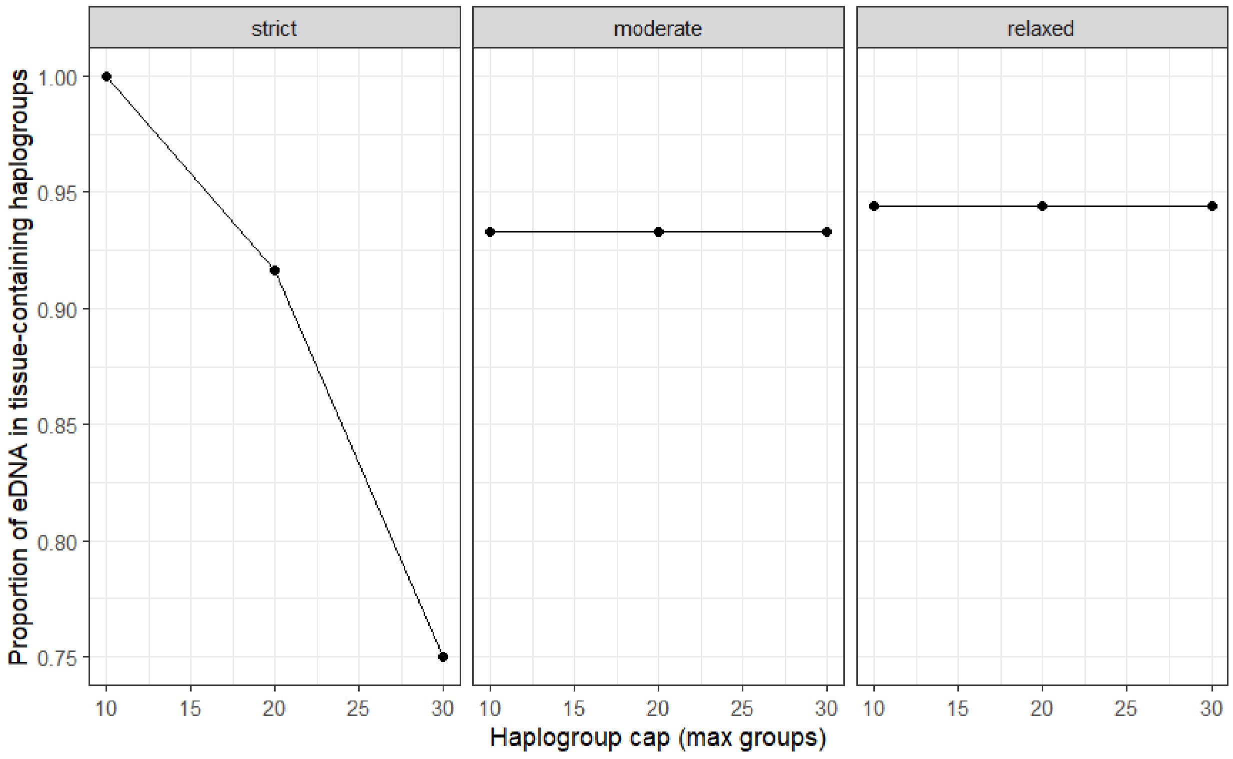


Supporting Figure 2. Sensitivity of haplogroup concordance between paired tissue and eDNA samples to the maximum number of allowed haplogroups (haplogroup cap) under strict, moderate, and relaxed filtering profiles. The y-axis shows the proportion of paired samples in which eDNA and tissue haplogroup assignments match. Concordance remains high under relaxed and moderate filtering across haplogroup caps, whereas strict filtering shows reduced concordance at higher caps.

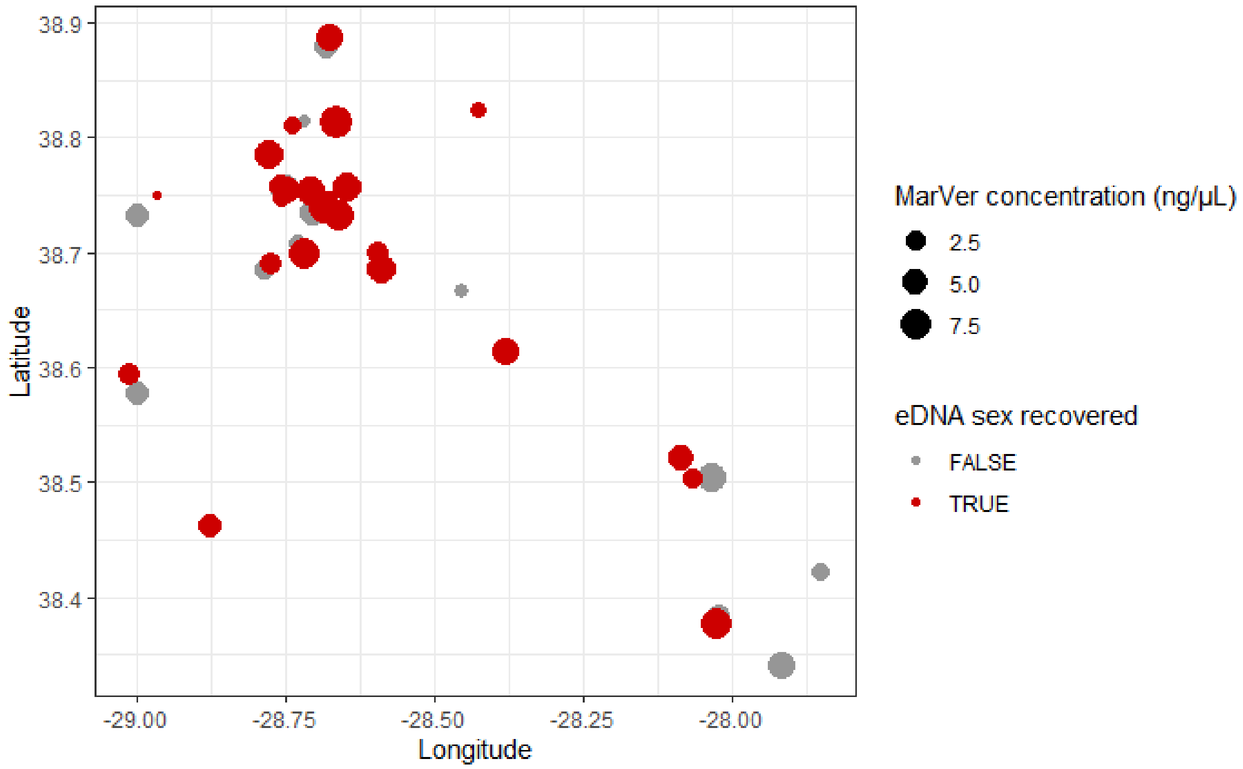


Supporting Figure 3. Spatial distribution of eDNA sampling locations included in sex determination analyses. Circle size corresponds to marine vertebrate (MarVer) DNA concentration (ng/µL) measured in each sample, and color indicates whether eDNA-based sex assignment was successfully recovered (red) or not (grey). Points represent individual sampling events.

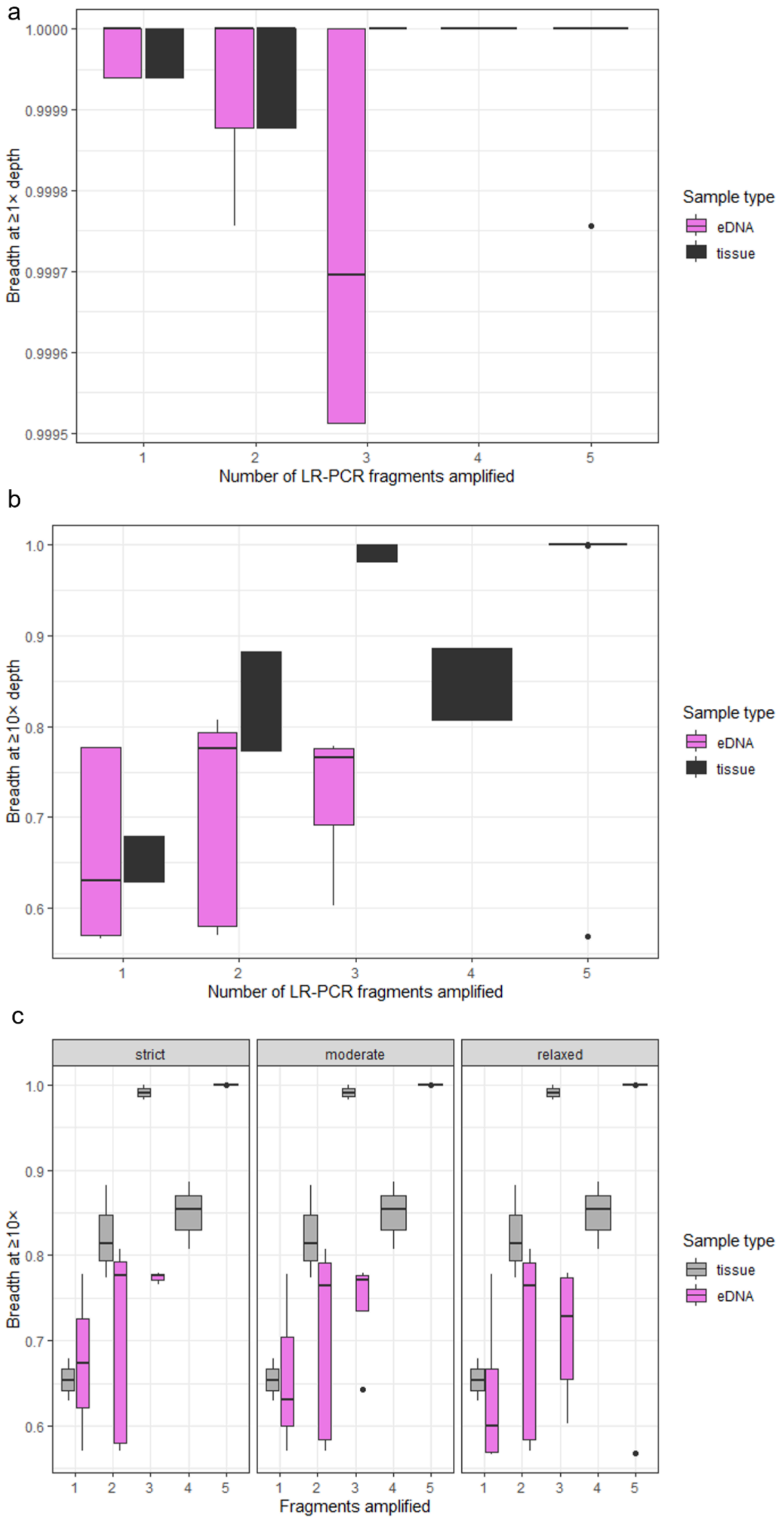


Supporting Figure 4. Relationship between the number of successfully amplified long-range PCR (LR-PCR) mitochondrial fragments and sequencing performance metrics for tissue and eDNA samples. Boxplots compare tissue (grey) and eDNA (purple) samples across increasing numbers of amplified fragments. Coverage breadth is calculated as the proportion of mitochondrial positions meeting the specified depth threshold. a) Breadth of mitogenome coverage at ≥1× depth. b) Breadth of mitogenome coverage at ≥10× depth. c) Breadth of coverage at ≥10× depth shown separately for strict, moderate, and relaxed bioinformatic filtering profiles.

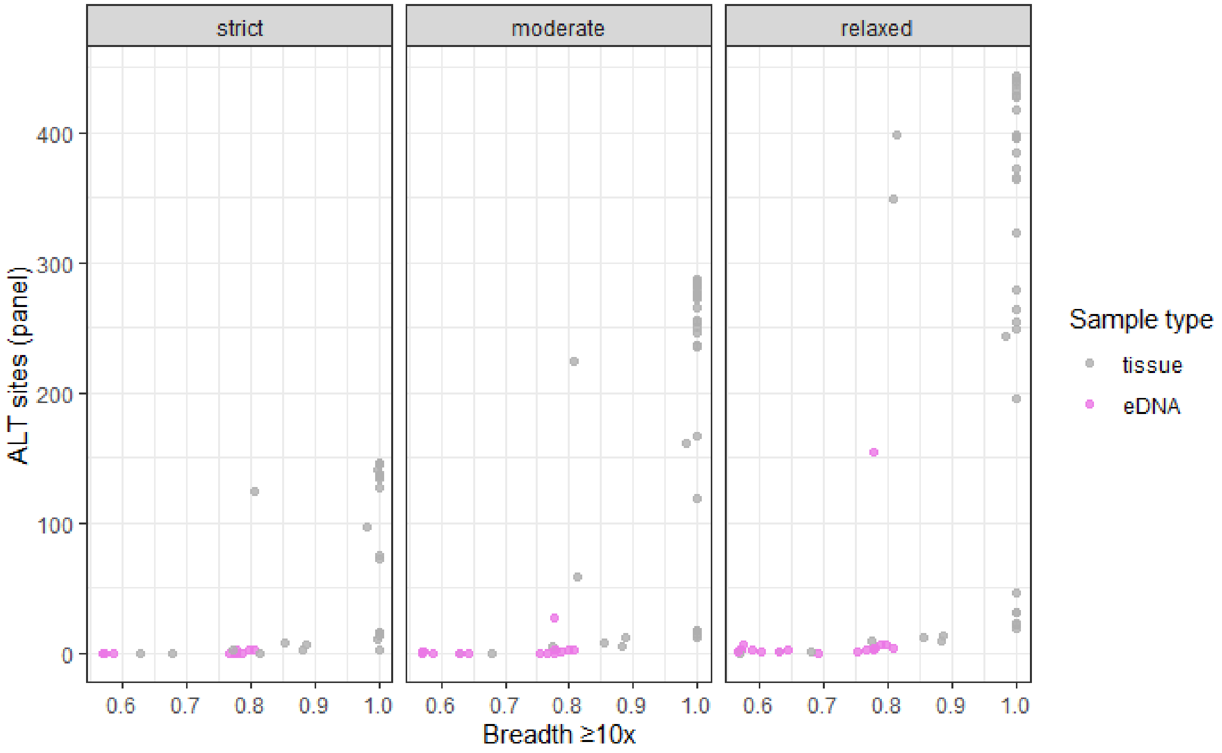


Supporting Figure 5. Relationship between mitogenome coverage breadth (≥10× depth) and the number of panel ALT sites detected per sample under strict, moderate, and relaxed bioinformatic filtering profiles. Points represent individual samples, colored by sample type (tissue, grey; eDNA, purple). Tissue samples exhibit substantially higher ALT site counts, particularly under relaxed filtering, whereas eDNA samples show consistently low ALT counts across coverage levels.

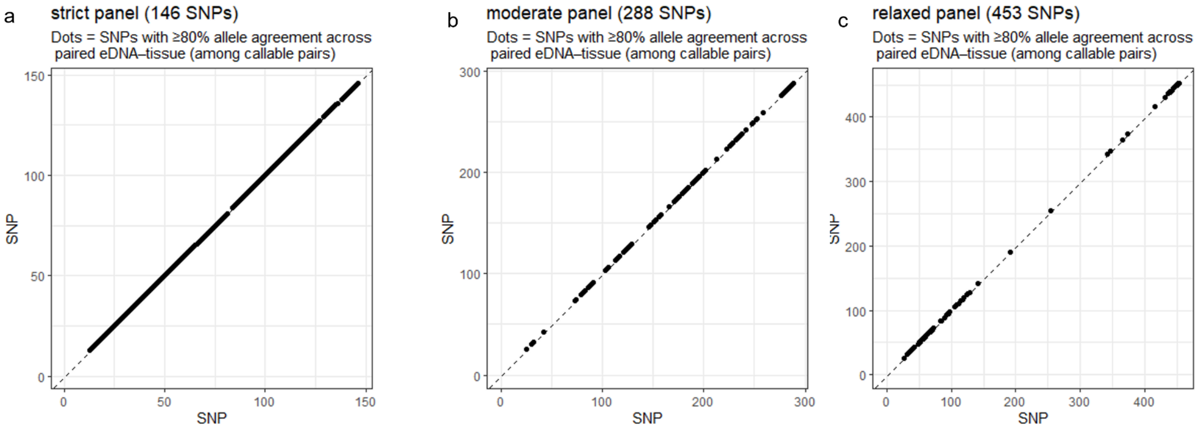


Supporting Figure 6. SNP-level allele concordance between paired tissue and eDNA samples across bioinformatic filtering profiles. Panels show a) strict (146 SNPs), b) moderate (288 SNPs), and c) relaxed (453 SNPs) SNP panels. Each point represents a SNP exhibiting >80% allele agreement between paired tissue and eDNA samples, with allele frequency in tissue plotted against the corresponding eDNA sample. The dashed diagonal indicates perfect (1:1) agreement.

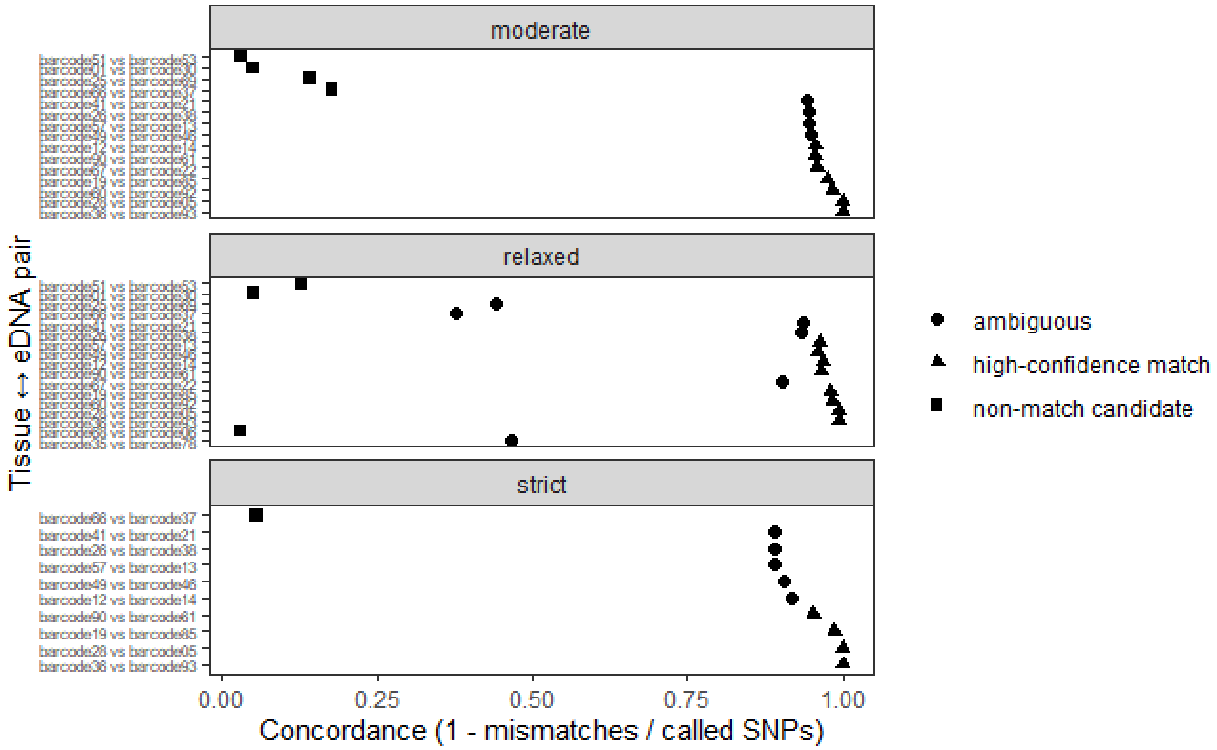


Supporting Figure 7. Pairwise SNP concordance between tissue and eDNA samples under strict, moderate, and relaxed filtering profiles. Concordance values are computed as 1 − (mismatches / called SNPs) for each potential tissue–eDNA pairing. Symbols denote classification of pair status (high-confidence match, ambiguous, or non-match). High-confidence matches consistently cluster near complete concordance across filtering profiles.

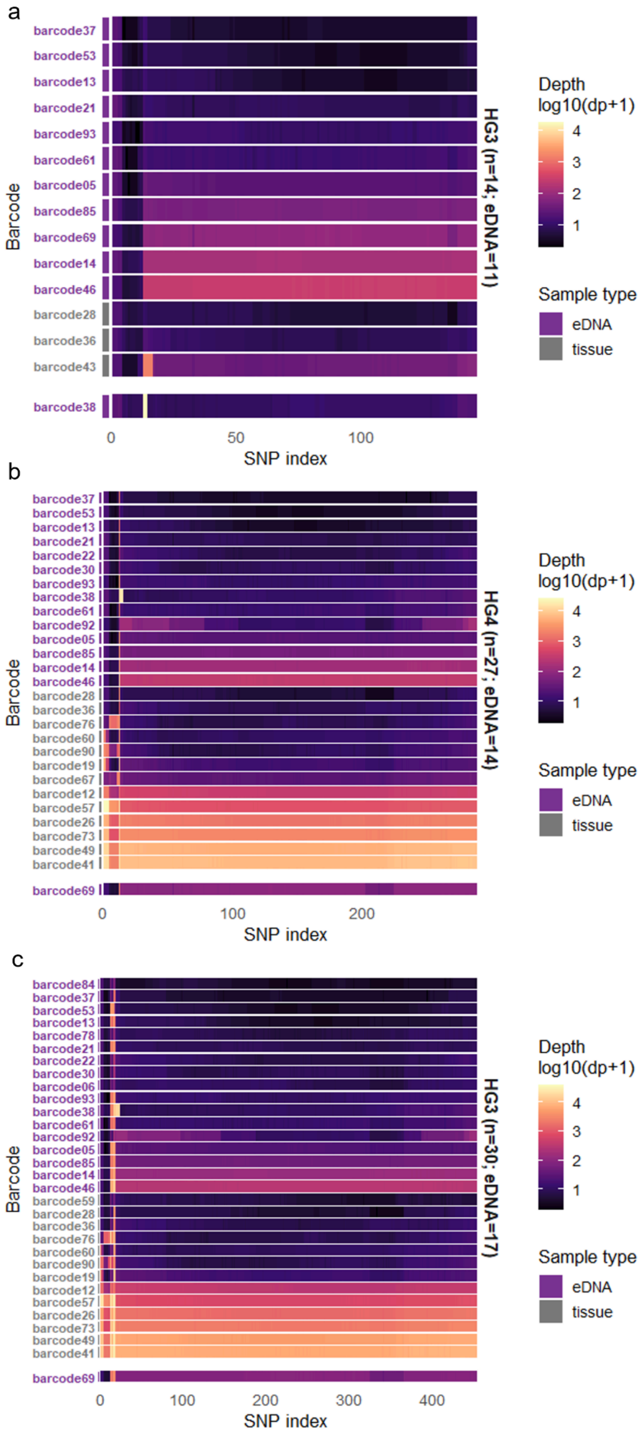


Supporting Figure 8. Depth structure across mitochondrial SNP positions (by haplogroup). Heatmaps show log-transformed sequencing depth across SNP positions for strict (a), moderate (b), and relaxed (c) filtering profiles. Rows represent individual sample barcode names (colored purple for eDNA, grey for tissue) and each location on the x-axis corresponds to a SNP position. Smaller haplogroups exhibit coherent depth profiles across SNP positions while the dominant haplogroup (HG4) exhibits greater heterogeneity in depth but consistent support across variants.

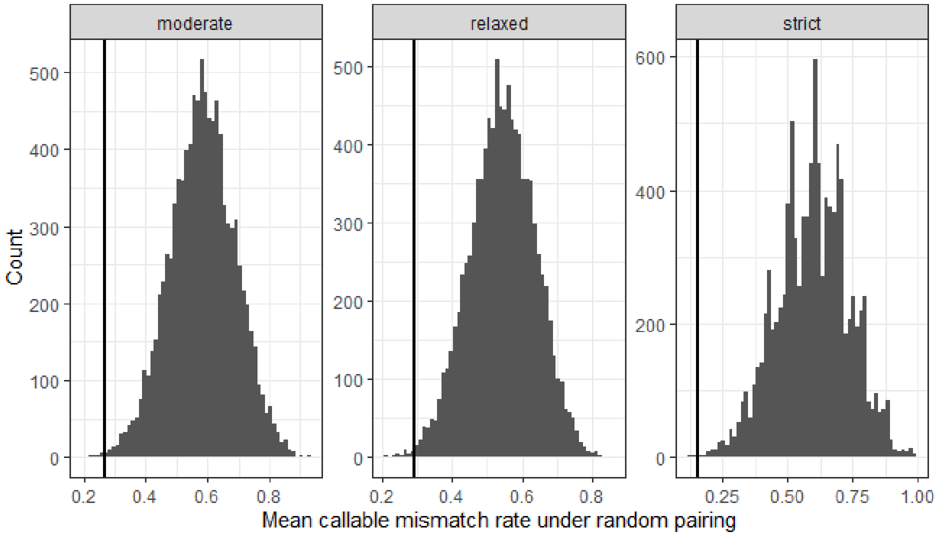


Supporting Figure 9. Null distributions of mean callable mismatch rate under random eDNA/tissue pairing across moderate, relaxed, and strict filtering profiles. Histograms represent the distribution of mean mismatch rates obtained from permutation tests in which tissue and eDNA samples were randomly paired. The vertical line indicates the observed mean mismatch rate for the true paired samples. Observed values falling outside the bulk of the null distribution indicate non-random concordance between paired samples.

Supporting Table 1. Primers used in this study for assessing the presence of vertebrate DNA, sperm whale-specific molecular sexing markers, and mitochondrial DNA haplotypes.

| Purpose | Reference | Sequence (5' - 3') | Fragment length (bp) | Target Gene |
|---|---|---|---|---|
| Vertebrate DNA Screening (MarVer3) | (Valsecchi et al., 2020) | F: MarVer3F AGACGAGAAGACCCTRTG<br>R: MarVer3R GGATTGCGCTGTTATCCC | 245 | 16S |
| Sexing | Modified from (Nishida et al., 2003) | F: Pm-SRY-F GTAGAAATAACCCTTGAATAGC<br>R: Pm-SRY-R CTCACCGTTCACAATTCTG | 127 | SRY |
| Sexing | This study | F: Pm-ZFX-F AAAAACCATCC+TGAACACCTT+A<br>R: Pm-ZFX-R +TTTACC+GCACTCCACACA+T | 336 | ZFX |
| Haplotyping Fragment 1 | (Alexander et al., 2013) | F: 1.4UPF AATCCAGGTCGGTTTCTATCT<br>R: Pma6916tSerR GTTCGAKTCCTTCCTTTCTT | 4,391 | 16S rRNA, ND1, ND2, COX1 |
| Haplotyping Fragment 2 | (Alexander et al., 2013) | F: Pma6800CO1F GAGAAGCMTTYRCATCCAAACG<br>R: PmaHDND4R GGGTCAGAGAAGAACGTAAGGG | 3,608 | COX1, COX2, ATP6, COX3, ND3, ND4L |
| Haplotyping Fragment 3 | (Alexander et al., 2013) | F: Mys10000ND4LF CGATCCCACCTAATRTCCGCA<br>R: Mys13000ND5R GCTCAGGCGTTGGTATAAGA | 3,024 | ND4L, ND4, ND5 |
| Haplotyping Fragment 4 | (Alexander et al., 2013) | F: PmaHS13660F GCCTYAACCAACCYTAYCTRG<br>R: Pma12sRNAR GTGCTTGATACCWGCTCCTTTT | 3,874 | ND5, ND6, CYTB, Control Region, 12s |
| Haplotyping Fragment 5 | (Alexander et al., 2013) | F: PmaM13tpheF_mod AAAGCAAGACACTGAAGATGTCT<br>R: PmaHD3106R_mod AACAGTCAGGCTGGATATTGC | 3,091 | 12S rRNA, 16S rRNA, ND1 |

F codes for "forward primer"; R codes for "reverse primer"; primer names are displayed as originally published.

Supporting Table 2. Distribution of pairwise mitochondrial SNP distances between matched eDNA/tissue samples.*

| profile | n SNPs perfect match | n SNPs < 10 | n SNPs < 20 | n SNPs < 50 | n SNPs < 100 | n SNPs < 200 | n SNPs < 300 | n SNPs < 400 |
|---|---|---|---|---|---|---|---|---|
| **relaxed (n = 453 SNPs)** | 0 | 4 | 8 | 11 | 11 | 11 | 14 | 17 |
| **Moderate (n = 288 SNPs)** | 2 | 4 | 11 | 11 | 11 | 11 | 15 | NA |
| **strict (n = 146 SNPs)** | 2 | 4 | 9 | 9 | 9 | 10 | NA | NA |

* For each profile, the number of paired samples falling below the increasing SNP distance thresholds is shared. Perfect match indicates zero SNP differences between paired samples and subsequent columns represent the counts of pairs differing by fewer than the specified number of SNPs.
